# Enhancing nasal retention: using the mucosal blanket to structure sprays

**DOI:** 10.64898/2026.01.12.698967

**Authors:** Julie Tzu-Wen Wang, Chi Wai Tsang, Nga Chi Yip, Shih-Che Hsu

## Abstract

The anatomy of the nasal cavity, although volumetrically relatively small, houses an effective surface area ca. 9.6 m^2^, which facilitates conditioning of air (humidity/temperature) through heat and moisture exchange using a dense vascularised network. Although effective at regulating air intake, the large surface area, permeable membranes and efficient transport systems consequently leave it vulnerable to airborne pathogens and high rates of infection. Recently, there has been a movement to improve nasal sprays, trying to maximise spray coverage whilst maintaining a viscous nature to extend residence across the mucosal lining. In this study, NoriZite®, a commercial nasal spray based on a natural polysaccharide, gellan, has been studied for its retention within healthy volunteers. Data collected shows elimination kinetics with a half-life of 4 hrs, remaining detectable in the nasal cavity for up to 6 hrs. As such, NoriZite® shows potential as a long lasting nasal barrier, which could be used for the prevention of airborne infections.

## Introduction

The nose plays two fundamental roles in human physiology, respiration and olfaction (sense of smell). As such, structurally it can be viewed as two primary areas, the olfactory region, located at the roof of the nasal cavity which contains a dense network of sensory neurons, and, the respiratory region, which is responsible for conditioning inhaled air.^1^ This unique anatomical setup, maximising surface area to volume ratio, is achieved through the nasal tubinates, curved, shelf-like structures that project into the nasal passages, facilitating the 150 cm^2^ surface whose primary role is to regulate airflow as it enters the body.^2^ Ultimately, this allows the two major function of the nose to be achieved: 1) climate control, adjusting air temperature and humidity, and; 2) filtration and defence, trapping of particles and pathogens.^3–5^ The latter, achieved through the mucosal lining which covers the nasal surfaces, a specialized tissue composed primarily of pseudostratified columnar epithelium formed of goblet cells and ciliated epithelial cells: not only resulting in an effective capture area of ca. 9.6m^2^, but producing a viscoelastic blanket which traps and removed foreign particles.^2,6,7^

The mucin blanket, a dynamic gel-like layer that overlays the nasal mucosa, is primarily composed of mucins; high-molecular-weight glycoproteins secreted by goblet cells and submucosal glands.^8–10^ These mucins form a viscoelastic matrix that traps inhaled particles, pathogens, and allergens, preventing them from reaching the epithelial surface.^11,12^ To achieve this action, the mucin blanket is organized into two distinct phases: a gel layer on top, which provides structural integrity and traps particulates, and a sol layer beneath, which allows for the coordinated movement of cilia.^13,14^ Ultimately, the ability to trap and flow at the same time ensures continuous renewal and removal of debris. Biologically, the mucin blanket contains not only mucins but also antimicrobial peptides, enzymes (such as lysozyme), immunoglobulins (especially IgA), and electrolytes, all of which contribute to its protective and immunological functions.^8–11^ The composition and thickness of the mucin layer can vary in response to environmental stimuli, infections, or inflammation, influencing both its barrier properties and its permeability. Moreover, the mucin blanket presents an opportunity, offering a targetable medium for mucoadhesive formulations designed to prolong residence time within the nasal cavity.^15–17^

The retention time of nasal sprays within the nasal cavity is a critical factor influencing drug absorption and therapeutic efficacy. Typically, conventional sprays exhibit short residence times, often clearing within 15 minutes.^18^ However, advanced formulations have significantly extended this duration. For instance, Bentrio nasal spray, a thixotropic gel-based formulation, demonstrated a residence time of up to 3.5 hours, maintaining a protective film across the nasal mucosa and oropharynx.^19^ Studies comparing various dosage forms – solutions, powders, gels, and mucoadhesive systems – show that mucoadhesive polymers can enhance retention.^20^ Such effectiveness is dependent on a range of factors including viscosity, spray pattern, droplet size, and device design, all of which influence deposition and clearance dynamics. These advances have been crucial for improving bioavailability, especially for drugs targeting systemic circulation or the central nervous system via the nasal route.^21–23^

To date, mechanical behaviour, in particular viscosity has been critical in determining retention within the nasal cavity. Most commercial nasal sprays exhibit shear-thinning behaviour, crucial for enabling atomization and droplet formation under the high shear rates (10^5^-10^6^ s^−1^) encountered when being pumped through an applicator.^24^ More advanced formulations, including bentonite-based gels, maximise the viscosity recovery post application, aiding resistance to mucociliary clearance and maintaining mucosal coverage for up to 3.5 hrs.^19^ Similarly, thermosensitive poloxamers have enabled in situ gelation, forming viscous solutions at body temperature.^25–27^ These “high” viscosity systems enhance retention, however, must be carefully balanced to avoid compromising sprayability and deposition. Recently, NoriZite® a gellan-based composite has demonstrated the ability to retain high viscosity, however, easily undergo atomisation within a manual spray applicator to achieve enhanced spray coverage.^28^

This work reports further development of NoriZite®, as a long term protective spray. The hypothesis, based on previous works showing that mucins found within the mucin blanket provide a structurally recognisable motifs, which act as a template for in situ structuring formation, will enable long term retention of the spray. A quantitative analytical technique, HPLC, was used to determine the presence of spray over n 8 hr timeframe and elimination kinetics compared to literature. To this end, an in vitro study was performed to establish a deeper understanding on mechanism for retention.

## Methods

NoriZite® Nasal Spray (Birmingham Biotech LTD., UK), Carrageenan (Sigma Life Science, UK), Gelatine (porcine, type A) (Sigma Life Science, UK), mucin (porcine stomach, type II) (Sigma Life Science, UK), PBS (Sigma Life Science, UK), Gellan gum (CG-LA) (CP Kelco), Type 1 water (Milli-Q, Merck Millipore).

### Nasal Spray retention in healthy volunteers

A healthy volunteer study was conducted by LabWide Solutions HK Limited (Hong Kong) using products registered by the local regulatory authorities, and in accordance with local ethical guidelines; strictly adhering to the principles outlined in the Belmont report (1979). In brief, healthy volunteers (n=5) were first swabbed to obtain baseline controls prior to the application of NoriZite® nasal spray. Following, volunteers applied two actuations of the spray to each nostril and were then instructed to sit quietly and gently breathe through their nose for 1 minute before resuming their normal daily routine. Swabs were obtained at 1 and 2 hrs, independently swabbing the left nostril and right, respectively. Subsequently, a minimum of 24 hrs was allowed for the nasal passage to regain homeostasis and the process repeated for 3 and 4 hrs. This was then repeated a third time for 6 and 8 hrs. Nasal swabs were flushed with 1 mL of HPLC grade water before discarding the cotton bud. Samples were then digested in equal amounts of HCl (2 M) in an autoclave (TOMY-SX-700) for 20 min at 121 °C (1bar). Once cooled, samples were neutralised with NaOH (2 M) and passed through a 0.22 mm filter. Calibration standards were prepared in the same manner, using 100 mL of NoriZite® in 900 mL of HPLC grade water. Post digestion and neutralisation, samples were serial diluted to create a 6-point calibration curve.

*HPLC* - analysis was conducted using a Thermo Separation SpectraSystem configured with an AS3000 auto sampler with column heater (set to 35 °C), Phenomenex Kinetex 5µm EVO C18 100Å, P4000 gradient pump, SCM1000 degasser and UV6000LP photodiode array detector (265 nm). Mobile phase “A” comprised of HPLC grade water with formic acid (1% v) and “B” HPLC grade methanol. 100 uL of sample was injected into the system and analysed at a constant flow rate of 0.5 mL.min^-1^, using a 2:1 (A:B) ratio for 15 min. The column was then washed using 100% HPLC grade methanol for 10 mins, before priming for a further 10 minutes.

## Results and discussion

Retention of the NoriZite® within the nasal cavity was studied via nasal swabbing and quantitative analysis with HPLC. Baseline controls were established by systematically digesting – high temperature acid (HCl (2 M)) treatment followed by neutralisation (NaOH (2 M)) – potential contaminates and pure NoriZite®. Chromatograms highlighted a capacity to deconvolute between the nasal contaminants, sampling swab and spray, where metabolites of the spray composite could be detected at ca. 11 mins (Figure 2a).

**Figure 1:**
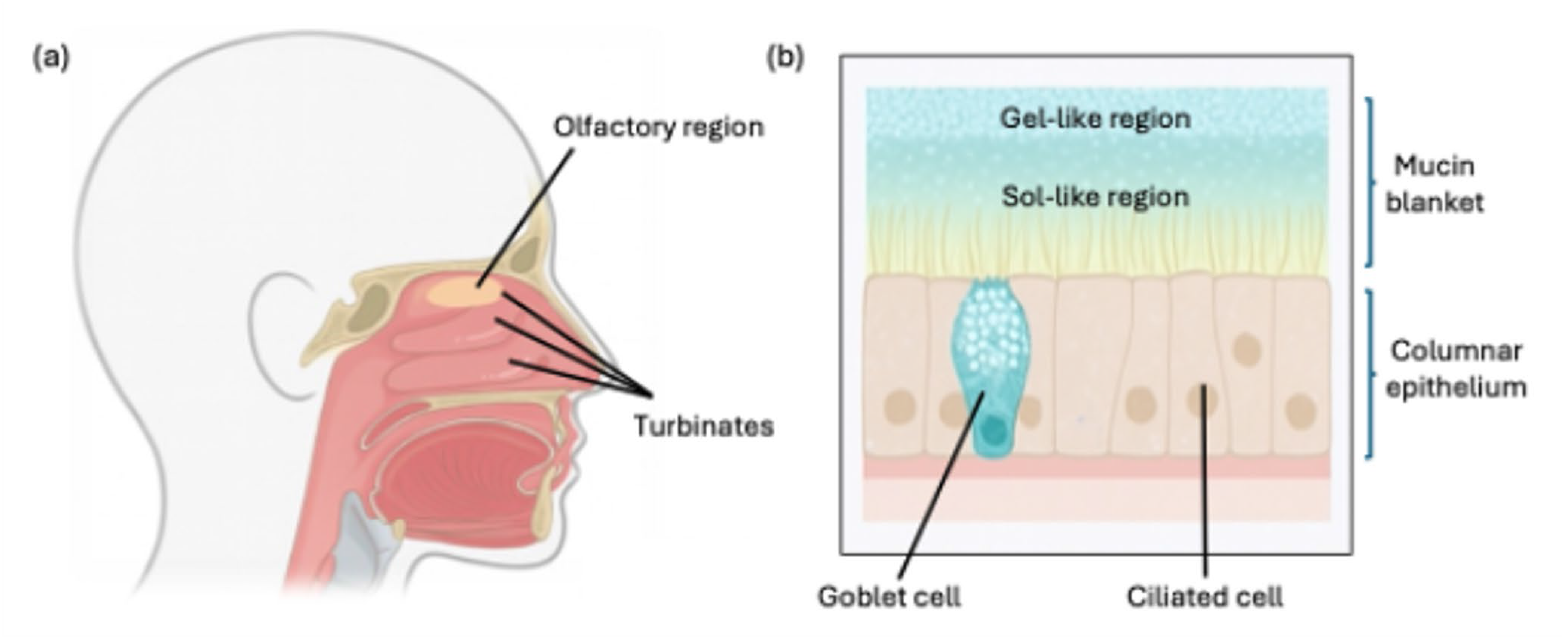
Anatomic images of the nasal cavity. a) schematic diagram of the nasal cavity highlighting turbinates and olfactory region. b) schematic representation of the mucosa highlighting cellular components, mucin blanket and identifying regions of gel- and sol-like mechanical behaviours.

**Figure 2:**
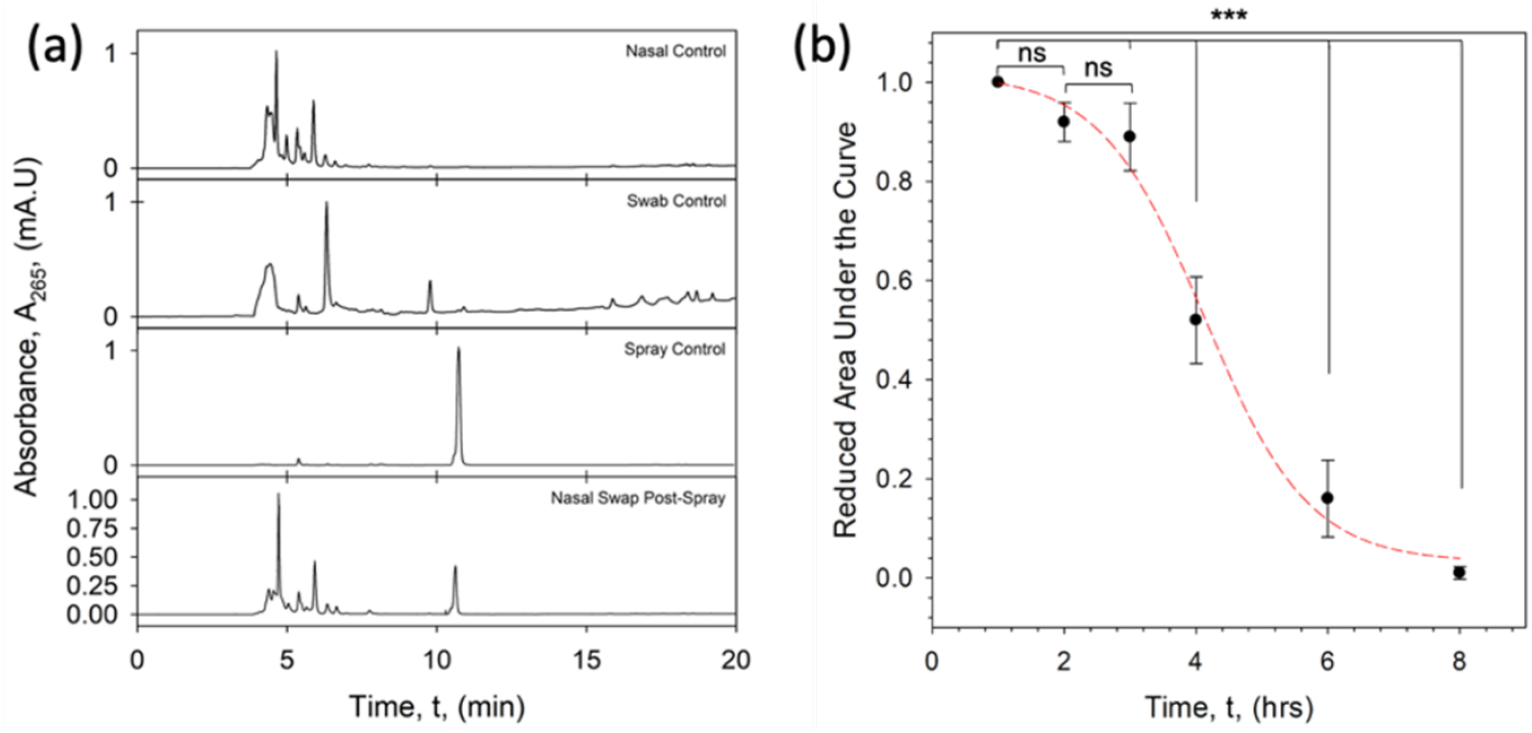
NoriZite® retention when applied to the nasal cavity of healthy volunteers (n=5). (a) HPLC data used to quantify amounts of nasal spray present at each timepoint. Plot shows chromatograms obtained for various controls used to deconvolute volunteer swab samples. (b) Elimination curve for the nasal spray detected in the nasal passages of volunteers over 8 hrs plotted as reduced values to account for variation between users. Data shows detection of the spray for up to 6 hrs.

Following establishment of baseline chromatograms, nasal swabs of healthy volunteers (n=5) were obtained over a time course of 8 hrs: hourly intervals, whereby each data point was obtained using a fresh application, ie. application followed by sampling, re-application 24 hrs later followed by sampling at the next required timepoint etc. To reduce inter-participant variability (changes in natural mucus levels, mucosa surface area, spray technique etc.), relative changes (*A*_*rel*_) in spray were obtained using eq1:

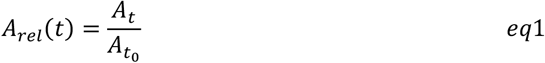

Following normalisation, values for the area under the curve (*A*) at each timepoint were averaged across all 5 participants and error determined as the 95% confidence interval. Figure 2b shows elimination kinetics for the spray, highlighting a sigmoidal response, as shown by the fit (*R*^2^ = 0.98) with first order behaviour following the primary plateau (determined by a significant decrease (p<0.001)). Moreover, the spray demonstrated a half-life (*t*_0.5_) of ca. 4 hrs, reaching the lower level of quantification (LLOQ) by 8 hrs.

The ability to slow nasal clearance using thickeners, for example hydroxypropyl methylcellulose (HPMC), has long been known.^29–31^ Indeed, labelled sprays with controlled thicknesses have been shown to express retention half-lives (*t*_0.5_) as a linear function of viscosity:^29^ low viscosity sprays (ca. 40 mPa.s) exhibiting complete clearance within 60 mins and those with higher viscosities (ca.400 mPa.s) demonstrating *t*_0.5_ much closer to 2 hrs – in healthy volunteers. Interestingly, the data presented here did not follow previous reports, maintaining a zero-shear viscosity of 200 mPa.s (data previously published),^28,32^ but with no significant clearance within the first 2 hours, once *in situ*. Moreover, the NoriZite® demonstrated *t*_0.5_ of 4 hrs remaining detectable within the nose for up to 6 hr following application (Figure 2b). Such differences have been attributed to the chemistry of the polymers used. Indeed, when in contact with a simulated mucosa, synergistic effects arising through mucin-gellan interactions have been reported, highlighting a time-dependent “stiffening” of the sprayed layer.^32^ It is proposed that the ability to structure across the mucin blanket indeed leads to such unexpected longevity.

## Conclusions

The ability to enhance the retention of nasal sprays across the mucosa has significant benefits across a range of pharmaceutical sectors. NoriZite® a gellan-based composite has demonstrated, in healthy volunteers, the ability to reside within the nasal cavity for up to 6 hrs. On comparison to literature, NoriZite® shows unexpectedly long retention for its viscosity. This suggests a mechanism which go beyond simple thickening, but the potential of synergistic behaviour between the native mucin blanket and spray chemistry. Ultimately, modification of mucocillary clearance enables NoriZite® to act as a barrier for extended timeframes.

